# Maximum CO_2_ diffusion inside leaves is limited by the scaling of cell size and genome size

**DOI:** 10.1101/2020.01.16.904458

**Authors:** Guillaume Théroux-Rancourt, Adam B. Roddy, J. Mason Earles, Matthew E. Gilbert, Maciej A. Zwieniecki, C. Kevin Boyce, Danny Tholen, Andrew J. McElrone, Kevin A. Simonin, Craig R. Brodersen

**Affiliations:** Institute of Botany, University of Life Sciences and Natural Resources, 1180 Vienna, Austria; School of Forestry & Environmental Studies, Yale University, New Haven CT 06511 USA; Department of Viticulture & Enology, University of California, Davis, CA 95616 USA; Department of Biological & Agricultural Engineering, University of California, Davis, CA 95616 USA; Department of Plant Sciences, University of California, Davis, CA 95616 USA; Department of Geological Sciences, Stanford University, Palo Alto, CA 94305 USA; USDA-Agricultural Research Service, Davis, CA 95616 USA; Department of Biology, San Francisco State University, San Francisco CA 94132 USA

**Author notes:** These authors contributed equally to this work (Guillaume Théroux-Rancourt, Adam B. Roddy).

## Abstract

Maintaining high rates of photosynthesis in leaves requires efficient movement of CO_2_ from the atmosphere to the chloroplasts inside the leaf where it is converted into sugar. Throughout the evolution of vascular plants, CO_2_ diffusion across the leaf surface was maximized by reducing the sizes of the guard cells that form stomatal pores in the leaf epidermis^1,2^. Once inside the leaf, CO_2_ must diffuse through the intercellular airspace and into the mesophyll cells where photosynthesis occurs^3,4^. However, the diffusive interface defined by the mesophyll cells and the airspace and its coordinated evolution with other leaf traits are not well described^5^. Here we show that among vascular plants variation in the total amount of mesophyll surface area per unit mesophyll volume is driven primarily by cell size, the lower limit of which is defined by genome size. The higher surface area enabled by smaller cells allows for more efficient CO_2_ diffusion into photosynthetic mesophyll cells. Our results demonstrate that genome downsizing among the flowering plants^6^ was critical to restructuring the entire pathway of CO_2_ diffusion, facilitating high rates of CO_2_ supply to the leaf mesophyll cells despite declining atmospheric CO_2_ levels during the Cretaceous.

## Main text

The primary limiting enzyme in photosynthesis, rubisco, functions poorly under low CO_2_ concentrations. For leaves to maintain high rates of photosynthesis, they must maintain high rates of CO_2_ supply from the atmosphere to the sites of carboxylation in the leaf mesophyll. The importance of maintaining efficient CO_2_ supply is reflected in the evolutionary history of leaf anatomy; leaf surface conductance has increased during periods of declining atmospheric CO_2_ concentration^2^, primarily due to increasing the density and reducing the sizes of stomatal guard cells through which CO_2_ diffuses^1,7,8^. However, allowing CO_2_ to diffuse into the leaf exposes the wet internal leaf surfaces to a dry atmosphere. Maintaining a high rate of CO_2_ uptake, therefore, requires high fluxes of water to be delivered throughout the leaf to replace water lost during transpiration (Figure S1), which is accomplished by a dense network of leaf veins^9,10^. Together, increases in the densities of leaf veins and stomata and reductions in guard cell sizes enabled the elevated photosynthetic rates that occurred only among angiosperm lineages throughout the Cretaceous despite declining atmospheric CO_2_ concentration^2,6,11–15^.

For a given leaf volume, the number of cells that can be packed into a space and the distance between different cell types is fundamentally limited by the size of these cells^15,16^. Because increasing investment in any one cell type will displace other cell types^17,18^, reducing cell size is a primary way of allowing more cell surface area of a given type to be packed into a given leaf volume. How small a cell can be is limited by the volume of its nucleus, which is commonly measured as genome size^19,20^. The reductions in cell size and increases in cell packing densities that occurred for veins and stomata only among angiosperm lineages during the Cretaceous, therefore, required reductions in genome size^6^. While reducing cell size and increasing cell packing density elevate maximum stomatal conductance to CO_2_^1,6^, realizing the potential benefits of elevated stomatal conductance would require modifications to the internal leaf structure that most limits CO_2_ transport: the absorptive mesophyll cell surface area exposed to the intercellular airspace.

Diffusion of CO_2_ inside the leaf is a major limitation to photosynthesis^3,4^ and has been considered for several decades to be a prime target for selection to increase photosynthetic capacity^21,22^. Unlike other tissues, the mesophyll is defined by both its cells and their surrounding intercellular airspace, both of which determine the overall CO_2_ conductance of the tissue. The conductance of the intercellular airspace (*g*_*ias*_) is thought to be much higher than the liquid phase conductance (*g*_*liq*_) of the cell walls and cell membranes because CO_2_ diffusivity is approximately 10,000 times higher in air than in water. These two conductances are arranged roughly in series, with *g*_*liq*_ acting as a greater limitation to CO_2_ uptake. While multiple factors, such as membrane permeability^4^, carbonic anhydrase^23^, and chloroplast positioning^24^ can be actively controlled over short timescales to regulate *g*_*liq*_, once a leaf is fully expanded, the structural determinants of *g*_*ias*_ and *g*_*liq*_, which include the sizes and configurations of cells and airspace in the mesophyll, are thought to be relatively fixed^4,22^. Of the various mesophyll traits commonly measured, the three-dimensional (3D) surface area of the mesophyll (*SA*_*mes*_) is thought to be the most important structural determinant of *g*_*liq*_. Because variation in leaf and mesophyll thicknesses influence *SA*_*mes*_ expressed on a leaf area basis^25^, standardizing *SA*_*mes*_ instead by tissue volume (*V*_*mes*_) accounts for variation in leaf construction^26,27^. The surface area of the mesophyll per tissue volume (*SA*_*mes*_*/V*_*mes*_; Figure S2), therefore, is the primary structural trait limiting CO_2_ diffusion from the intercellular airspace into the hydrated cell walls of the mesophyll. Because smaller cells have a higher surface area per volume than larger cells, we hypothesized that reducing cell size by genome downsizing would allow for higher *SA*_*mes*_*/V*_*mes*_ that results in higher rates of CO_2_ supply to the chloroplasts.

We tested this hypothesis using high resolution, three-dimensional (3D) X-ray microcomputed tomography (microCT) to characterize cell sizes, cell packing densities, and the exposed three-dimensional surface area of the mesophyll tissue of leaves spanning the extant diversity of vascular plants. The mesophyll tissue of most leaves is composed of two distinct layers, the palisade and the spongy mesophyll, that are thought to be optimized for different functions^28,29^. We analyzed these two layers separately to determine how differences in their 3D tissue structure (Figures S1 and S2) may drive differences in *g*_*ias*_ and *g*_*liq*_. We predicted, therefore, that in addition to heightened densities of veins and stomata, increasing *SA*_*mes*_*/V*_*mes*_ was an essential innovation unique to the angiosperms that enabled their elevated rates of CO_2_ supply to the mesophyll cells despite declining atmospheric CO_2_ concentrations during the Cretaceous^2,6,11–14,20^.

### Genome downsizing enables re-organization of the leaf mesophyll

We quantified from microCT images the sizes of spongy and palisade mesophyll cells and stomatal guard cells, as well as the packing densities of veins and stomata per unit leaf area to determine whether genome size limits the sizes and packing densities of cells in all leaf tissues for 87 species spanning the extant diversity of vascular. Genome size was a strong predictor of cell size in both the upper palisade mesophyll layer and the lower spongy mesophyll layer even after accounting for shared evolutionary history (Figure 1b-c; Supplementary Table S1), consistent with results for veins and stomata^6^ (Figures 1d, S3). As was expected, genome size did not correlate with porosity (Figure S3) because porosity is quantified as a volumetric fraction, and cell size can vary independently of porosity. Despite the role of porosity in facilitating diffusion in the intercellular airspace^30^, traits other than porosity related to cellular organization within the mesophyll are likely to have a greater influence on the diffusive conductance of CO_2_ through the intercellular airspace and into the photosynthetic mesophyll cells^27^.

**Figure 1.**
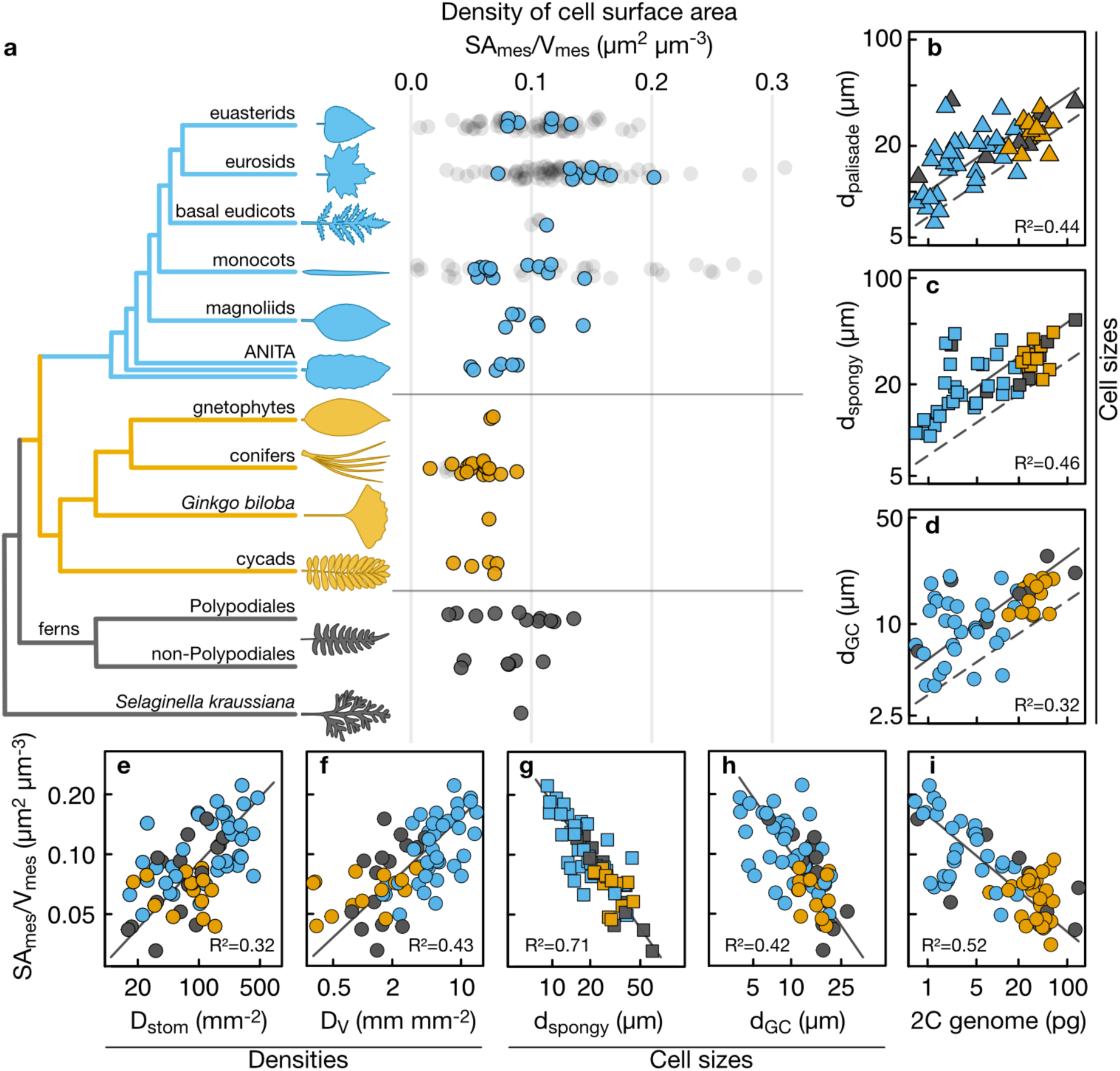
Mesophyll surface area per mesophyll volume (SA_mes_/V_mes_) scales with cell size, cell packing densities, and 2C genome size across vascular plants. (a) Distribution of SA_mes_/V_mes_ across 87 species of terrestrial vascular plants from all major clades (colored points) and compared to values computed from the literature (shaded dots, 81 angiosperms and four gymnosperms). The diameters of (b) palisade mesophyll cells (d_palisade_), (c) spongy mesophyll cells (d_spongy_), and (d) stomatal guard cells (d_GC_) are strongly predicted by genome size. SA_mes_/V_mes_ is positively related to the densities of (e) stomata on the leaf surface (D_stom_) and (f) veins in the leaf (D_V_). SA_mes_/V_mes_ is negatively related to (g) spongy mesophyll cellular diameter, (h) guard cell diameter, and (i) 2C genome size. Solid lines represent standardized major axes, and dashed lines represent the published relationship between meristematic cell volume and 2C genome size. All bivariate relationships remained highly significant after accounting for shared evolutionary history (Supplementary Table S1).

We tested whether the effect of genome size extends beyond limiting the sizes and packing densities of cells to influencing the surface area of the mesophyll tissue exposed to the intercellular airspace. We combined new measurements of *SA*_*mes*_*/V*_*mes*_ on the species for which we had microCT images with data extracted from the literature for 85 species (Figure 1a). *SA*_*mes*_*/V*_*mes*_ was coordinated with the packing densities of veins and stomata (Figure 1e,f), the diameters of spongy (Figure 1g) and palisade mesophyll cells (Supplementary Table S1), and the diameters of stomatal guard cells (Figure 1h). Furthermore, genome size was a strong predictor of *SA*_*mes*_*/V*_*mes*_ (Figure 1i). While small genomes, small cells, and high *SA*_*mes*_*/V*_*mes*_ occur predominantly among the angiosperms, some xerophytic ferns, as well as the lycophyte *Selaginella kraussiana*, also share these traits. The repeated co-occurrence of these traits among different clades (Supplementary Table S1) further corroborates the role of genome size in determining the sizes and arrangement of cells and tissues throughout the leaf that enable high fluxes of CO_2_ and H_2_O exchange with the atmosphere.

### Increasing liquid phase conductance optimizes the entire diffusive pathway

While light is intercepted primarily by the upper palisade mesophyll layer, CO_2_ enters the leaf on the lower spongy mesophyll layer for most terrestrial plants, creating opposing gradients of two of the primary reactants in photosynthesis. Within a leaf, the spongy and palisade layers have divergent cell shapes and organizations that are thought to accommodate these opposing gradients by facilitating CO_2_ diffusion in the gaseous and liquid phases. Both cell size and porosity can affect *SA*_*mes*_*/V*_*mes*_ and the diffusive conductances (*g*_*ias*_ and *g*_*liq*_) considered targets of selection to increase photosynthesis^20,25,30–32^. To determine whether cell size or porosity has a greater effect on *SA*_*mes*_*/V*_*mes*_, *g*_*ias*_, and *g*_*liq*_, we first quantified cell size, porosity, and *SA*_*mes*_*/V*_*mes*_ for the spongy and palisade layers separately for 47 species in our dataset, encompassing all major lineages of vascular plants.

The scaling of cell size with *SA*_*mes*_*/V*_*mes*_ (Figure 1g-h) suggested that cell size would have a greater impact than porosity on *SA*_*mes*_*/V*_*mes*_. Smaller cells have a higher ratio of surface area to volume, an effect that could propagate up to influencing *SA*_*mes*_*/V*_*mes*_ of the entire tissue. In contrast, we predicted that porosity would not have a consistent impact on *SA*_*mes*_*/V*_*mes*_ because at very low porosities there is very little cell surface area exposed to the airspace while at very high porosities there is very little cell surface area relative to a large volume of tissue. Consistent with these predictions, decreasing cell size led to higher *SA*_*mes*_*/V*_*mes*_ across species and mesophyll layers, and variation in porosity had no consistent effect on *SA*_*mes*_*/V*_*mes*_ (Figure 2). Rather, both low (<0.1) and high (>0.6) porosities led to lower *SA*_*mes*_*/V*_*mes*_. This conditional effect of porosity on *SA*_*mes*_*/V*_*mes*_ suggests that there is a relatively narrow range of porosities that allows for simultaneous optimization of *g*_*liq*_ and *g*_*ias*_. However, the strong and consistent effect of reducing cell size on increasing *SA*_*mes*_*/V*_*mes*_ among species and among mesophyll tissues within a leaf further implicates cell size and, by extension, genome size in controlling cell- and tissue-level traits responsible for increasing the CO_2_ conductance of the mesophyll.

**Figure 2.**
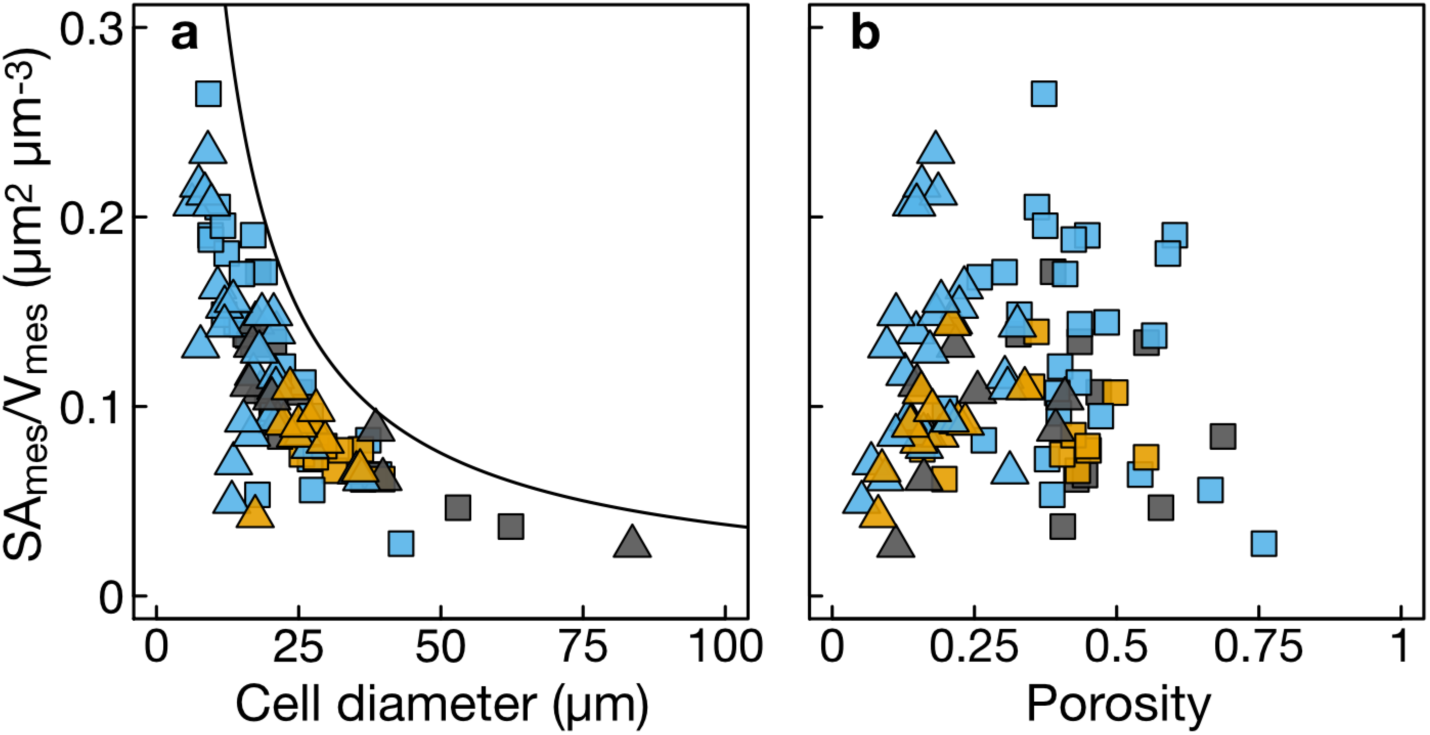
The effects of cell size and porosity on 3D mesophyll surface per mesophyll volume (SA_mes_/V_mes_). (a) Smaller cells in both the palisade (triangles) and spongy (squares) mesophyll are associated with higher SA_mes_/V_mes_. The solid line represents the theoretical maximum SA_mes_/V_mes_ calculated from the densest packing of cylinders of varying diameter in a rectangular volume (with a porosity of approximately 0.09 m3 m-3). (b) SA_mes_/V_mes_ was highest for leaves and tissues of intermediate porosity because the highest possible porosity can occur only when there are no cells and the lowest porosity occurs when all cells are in complete contact and there is no airspace. Points are colored by plant clade, according to Figure 1.

To test how these anatomical traits affect *g*_*ias*_ and *g*_*liq*_, we compared modeled estimates of *g*_*ias*_ and *g*_*liq*_ per unit leaf volume^4,27^ in which cell size and porosity were varied independently to anatomical measurements of the two mesophyll layers. Although this modeling did not incorporate adjustments that can alter *g*_*liq*_ over short timescales, it nonetheless shows how variation in anatomy, which is relatively fixed once a leaf has expanded^4^, can influence *g*_*ias*_ and *g*_*liq*_. Based on simple packing of capsules, we predicted that increasing volumetric *g*_*liq*_ would occur primarily by decreasing cell size, while increasing volumetric *g*_*ias*_ would occur primarily by increasing porosity. We also predicted that the palisade layer, whose densely packed columnar cells channel light deep into the leaf much as a fiber optic cable directs light^28^, would be optimized for *g*_*liq*_ rather than for *g*_*ias*_ in order to deliver CO_2_ efficiently to the places where light is abundant. In contrast, we predicted that the spongy mesophyll layer would be optimized for high *g*_*ias*_ in order to promote gaseous CO_2_ diffusion into the upper palisade layer^3^ while also scattering and absorbing light^33,34^.

Our modeling confirmed that cell size and porosity have different effects on volumetric estimates of *g*_*liq*_ and *g*_*ias*_ (Figure 3). While increasing porosity leads to higher *g*_*ias*_, it has a relatively small effect on *g*_*liq*_ for a given cell size. In contrast, increasing *g*_*liq*_ predominantly occurs by reducing cell size, which has only a moderate effect on *g*_*ias*_ and only when porosity is relatively high. Additionally, for a given cell size, increasing porosity reduces *g*_*liq*_. Thus, reductions in cell size increase both *g*_*liq*_ and *g*_*ias*_, but increasing porosity has opposite effects on *g*_*liq*_ and *g*_*ias*_. As predicted, the palisade layer had lower porosities that are associated with higher *g*_*liq*_, while the spongy layer had higher porosities that are associated with higher *g*_*ias*_ (Figures 3). This differentiation of function between the two layers reflects the need to maintain a high *g*_*ias*_ in the spongy mesophyll where CO_2_ is abundant to promote its diffusion into the palisade and the need to maintain high *g*_*liq*_ in the palisade mesophyll where light is abundant to promote liquid-phase diffusion of CO_2_ through the cell walls and into the chloroplasts. Many species, particularly angiosperms, have palisade mesophyll characterized by small, highly packed cells that allow volumetric *g*_*liq*_ to be higher than *g*_*ias*_ of this tissue (Figure 3). This pattern suggests that CO_2_ fixation in the palisade may be limited by the gaseous supply of CO_2_ and not by its liquid-phase diffusion into cells, consistent with prior reports for hypostomatous leaves that the majority of CO_2_ fixation occurs in the deeper palisade and not at the top of the leaf where CO_2_ is unlikely to penetrate^33,34^. The structure and organization of palisade and spongy layers of the mesophyll therefore reflect the relative strengths of the opposing gradients of CO_2_ and light.

**Figure 3.**
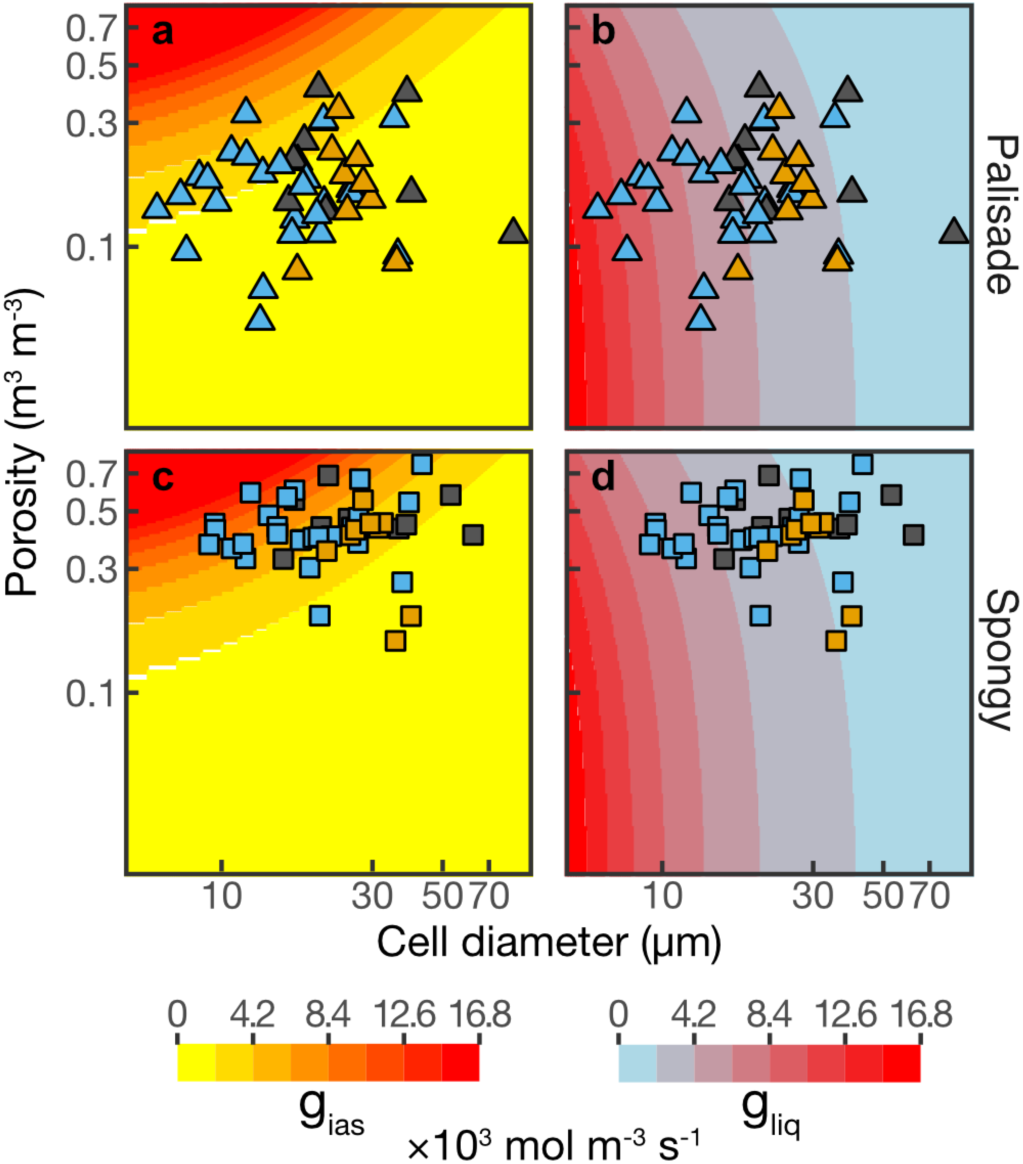
Distribution of observed mesophyll cell sizes and porosities relative to modeled estimates of gas phase conductance (g_ias_) and liquid phase conductance (g_liq_) to CO_2_. Measured values of cell size and porosity for the palisade (a-b, triangles) and spongy (c-d, squares) mesophyll layers are plotted over theoretical airspace conductance (g_ias_; a,c) and liquid phase conductance (g_liq_; b,d). Colored backgrounds represent conductances estimated from simulated leaves of varying cell diameter and porosity (see Supplemental Methods for details). Points are colored by plant clade, according to Figure 1.

## Conclusion

Our results suggest that the heightened rates of leaf-level gas exchange that occurred among Cretaceous angiosperms were coordinated with changes not only in veins and stomata^2,6,11,12,15,35^ but also in the three-dimensional organization of the leaf mesophyll tissues that limits the exchange of CO_2_ and water. Although coordinating changes in veins, stomata, and the mesophyll undoubtedly involves multiple molecular developmental programs, the scaling of genome size and cell size emerged as the predominant factor driving the increase in *SA*_*mes*_*/V*_*mes*_ and *g*_*liq*_ that together enabled higher rates of CO_2_ movement into the photosynthetic mesophyll cells. Because photosynthetic metabolism is the primary source of energy and matter for the biosphere, leaf-level processes are directly linked to ecological processes globally^8^. Yet, theory linking ecosystem processes to organismal level metabolism has focused predominantly on the structure of vascular supply networks^36,37^. Our results suggest that the scaling of photosynthetic metabolism with resource supply networks extends beyond the vascular system and into the photosynthetic cells of the leaf mesophyll where energy and matter are exchanged. Moreover, these results highlight the critical role of cell size in defining maximum rates of leaf gas exchange^6,20^, in contrast to assumptions in current theory that terminal metabolic units are size-invariant^38,39^. Incorporating the structure of the mesophyll tissue into theory linking leaf-level and ecosystem-level processes could improve model predictions of photosynthesis. Furthermore, the physiological benefits of small cells may be one reason why the angiosperms so readily undergo genome size reductions subsequent to genome duplications^6,20,40,41^. While whole genome duplications may drive ecological and evolutionary innovation^42–44^, selection for increased photosynthetic capacity subsequent to genome duplication may drive reductions in both cell size and genome size to optimize carbon fixation, reiterating a role for metabolism in genome size evolution.

## Methods

### Plant material

Fully expanded and mature leaves from healthy and well-watered plants were collected from various greenhouses, botanical gardens, fields, and other outdoor growing locations to represent a broad diversity of plant groups. Only C3 plants were used in this dataset. Leaves were cut at the base of the petiole or short stem segment, the cut end was wrapped in wet paper towels, and the entire shoot immediately put in a plastic bag before being transported to the synchrotron and scanned within 36 h of excision.

### microCT data acquisition

MicroCT scanning was carried out at the Advanced Light Source (ALS; beamline 8.3.2) at the Lawrence Berkeley National Lab (LBNL, Berkeley, CA, USA), the Swiss Light Source (SLS; TOMCAT Tomography beamline) at the Paul Scherrer Institute (Villigen, Switzerland), and the Advanced Photon Source (APS; beamline 2-BM-A,B) at Argonne National Laboratory (ANL, Lemont, IL, USA). Samples were prepared before each scan (less than 30 min) and, for laminar leaves, a ∼1.5 to 2-mm-wide and ∼15-mm-long piece of leaf tissue was excised between the mid-rib and the leaf outer edge. For needle and non-laminar leaves, a piece of leaf ∼15-mm-long was cut out. Tissue samples were then enclosed between two pieces of Kapton (polyimide) tape to prevent desiccation while allowing high X-ray transmittance. They were mounted in the sample holder, centered in the microCT X-ray beam, and scanned using the continuous tomography mode capturing 1,025 (ALS, APS) or 1,800 (SLS) projection images at 21 to 25 keV, using a 5x, 10x, or 40x objective lens, yielding a final pixel resolution between 1.277 - 0.1625 µm. Each scan was completed in 5 min to 15 min.

Image reconstruction was carried out using TomoPy^45^, a Python-based framework for reconstructing tomographic data, for all ALS samples, or using the in-house reconstruction platform for SLS or APS samples. Each leaf scan was reconstructed using both the gridrec^46^ and phase retrieval^47^ reconstruction methods, except for APS samples which possessed only gridrec reconstructions. Image stacks were cropped to remove tissue that was dehydrated, damaged, or contained artifacts from the imaging or reconstruction steps. Laminar leaves were aligned so that the epidermis was parallel to the image canvas top and bottom border. The final stacks contained ∼500-2000 eight-bit grayscale images (downsampled from 16 or 32-bit images).

### Leaf trait analysis

Leaf and mesophyll thickness were measured on the final gridrec image stack in cross sectional view. Cell diameter was measured on a minimum of 10 cells for each of the palisade and spongy layers on the gridrec image stack in paradermal view, as well as for the guard cell length and diameter. For spongy cells with trilobed or irregular shape, cell diameter was measured on the arms of the cells and not at their center.

Stomatal density and vein density were measured on the original, i.e. uncropped, image stack to capture the largest surface area by counting the number of stomata or measuring the length of veins present.

To extract surface area and volumes, the final image stacks were first segmented using manual or automated methods of Théroux-Rancourt et al.^26,48^ to segment the mesophyll cells, the airspace, the vasculature (combined veins and bundle sheath), and the background (including the epidermis). For both methods, ImageJ^49^ was used to segment or prepare the image stacks for segmentation. Airspace (*V*_*pores*_), mesophyll cell (*V*_*cells*_), and vasculature volume (*V*_*veins*_), as well as the surface area exposed to the intercellular airspace (*SA*_*mes*_) were then extracted using methods from Théroux-Rancourt et al.^26^ with the ImageJ plugin BoneJ^50^, or using a custom Python program^48^ (https://github.com/plant-microct-tools/leaf-traits-microct). The total mesophyll volume (*V*_*mes*_) was computed as the sum of *V*_*pores*_ and *V*_*cells*_. The area of the image stack in paradermal view was used as the leaf area (*A*_*leaf*_) of the image stack and used to compute S_m_, the exposed surface area per leaf area (*SA*_*mes*_*/A*_*leaf*_).

For the palisade and spongy data, stacks from within both tissues were cropped out of the segmented stack so as to capture a highly representative volume from these tissues, and involved cropping at the interface between both tissues, or where vasculature were highly present, to keep an accurate representation of surface area and cell diameter. Surface area and volumes (*V*_*pores*_ and *V*_*cells*_) were extracted using BoneJ.

### Genome size data

For the majority of our dataset, we matched existing 2C genome size (pg) data available in the Kew Plant DNA C-values Database^51^ (https://cvalues.science.kew.org). For several species in our dataset that were not available in the database, fresh leaf samples were collected from the same plants imaged using microCT from the University of California Botanical Garden, Berkeley CA. Genome sizes for these species were measured by the Benaroya Research Institute, Virginia Mason University, using the *Zea mays* or *Vicia faba* standards and following standard protocols^52^.

The relationship between meristematic cell diameter (*d*_*meristem*_) and genome size was computed based on the published relationship between genome size and meristematic cell volume for angiosperms^19^. Assuming that meristematic cells were spheres the cell diameter could be calculated as:

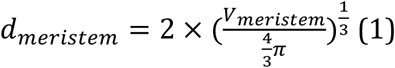

Deviations in cell shape from a sphere would allow for smaller diameters in one axis while maintaining the same cell volume.

### Phylogenetic analyses

To determine the evolutionary coordination between traits, we constructed a phylogeny from the list of taxa using Phylomatic (v. 3) and its stored family-level supertree (v. R20120829) using the R package brranching^53^. Following published methods^6^, we compiled node ages of named crown groups from fossil-calibrated estimates of crown group ages^54–56^. Of the 80 internal nodes in our phylogeny, 34 of them had published ages, which were assigned to nodes, and then branch lengths between dated nodes smoothed using the function ‘bladj’ in the software Phylocom (v. 4.2)^57^.

We tested whether there was correlated evolution between traits using phylogenetic least squares regression with a Brownian motion correlation structure, using the R packages nlme^58^ and ape^59^. For analyses of correlated trait evolution, traits were log-transformed to improve normality prior to regression analyses.

### Simulating conductance data using cell size and porosity

To simulate the range of liquid and intercellular airspace conductance used to generate the background of Figure 3, we used all possible combinations of cell diameter (0.1 µm steps) and porosity (0.01 steps) between 5 and 124 µm (1 µm below and 40 µm above the value range in our dataset) and 0.02 and 0.96 of porosity (0.03 below and 0.01 above tissue specific values in our dataset).

For the liquid phase conductance, *g*_*liq*_, we approximated cells as capsule shaped, as capsules give the best approximation of cell shape^60,61^. The capsule height was two times its diameter. We then generated the densest lattice possible, consisting of 30 cells in a (5 × diameter)^2^ projected area (Figure 4 below), giving a porosity of 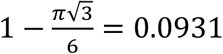 in the capsule body section, and ∼0.372 in the hemispheric ends (volume of a rectangular prism of *d* height and 5 *d* edges − volume of 30 spheres of 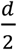 radius), for a total pore volume ratio of 0.186. The volume of that mathematically densest lattice, including cells and pores, equaled 2 × diameter × projected area. In cases where the simulated porosity was larger than 0.186, additional pore volume was added and so increased the total lattice volume, but not the cell volume. In cases where pore volume was below 0.186, pore volume was subtracted which decreased the total volume, but not the cell volume. This case represents when cells are inflated and deformed into each other, thus keeping similar surface area and volume, but reducing surrounding pore space.

**Figure 4.**
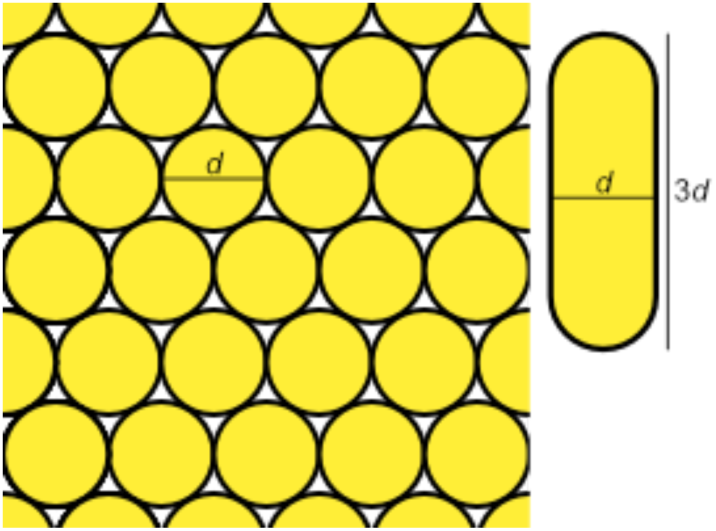
Schematic of the cell lattice used to simulate the intercellular airspace and liquid phase conductances. The lattice consisted of 30 capsule shaped cells of d diameter within a (5d)^2^ projected area, and 3d height, with a 2d length of the cylindrical body, for a total volume of 3d × (5d)^2^.

Liquid phase conductance per mesophyll volume was computed as in Evans et al.^4^ as a function of the surface area exposed to the intercellular airspace per volume, itself a function of cell diameter and porosity within the cell lattice:

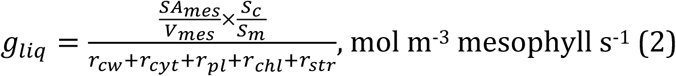

where *S*_*c*_*/S*_*m*_ is the proportion of cell surface occupied by chloroplasts (m^2^ chloroplast m^-2^ mesophyll surface area), assumed to be 0.85, and the different resistance (*r*_*i*_) components of the liquid diffusion path: *r*_*cw*_, cell wall; *r*_*cyt*_, cytosol; *r*_*pl*_, plasma membrane; *r*_*chl*_, chloroplast envelope; and *r*_*str*_, stroma. All resistances are assumed to be fixed and follow the average values used in Théroux-Rancourt and Gilbert^62^ for *r*_*cw*_ (10 m^2^ chloroplast s mol^-1^) and *r*_*cyt*_ (2 m^2^ chloroplast s mol^-1^), or used average values from Evans et al.^4^ for *r*_*pl*_ (7.1 m^2^ chloroplast s mol^-1^), *r*_*chl*_ (14.2 m^2^ chloroplast s mol^-1^), and *r*_*stroma*_ (7.1 m^2^ chloroplast s mol^-1^).

For intercellular airspace conductance, *g*_*ias*_, we used the equation of Earles et al.^27^, accounting for tortuosity (*τ*) and diffusive path lengthening from the stomata (*λ*) as functions of porosity (*θ*):

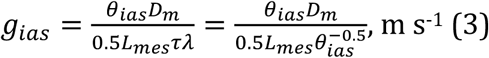

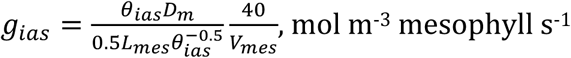

where *θ*_*ias*_ is the porosity of the mesophyll, *D*_*m*_ is the diffusivity of CO_2_ in the air (1.51×10^−5^ m^2^ s^−1^), and 0.5 *L*_*mes*_ is half the mesophyll thickness, i.e. half the cell lattice height. To convert to molar units, we used the conversion^63^ of 40 mol m^−2^ s^−1^ per m s^−1^ and divided by the lattice volume (*V*_*mes*_). Mesophyll thickness was considered to increase with an increase in cell diameter (*L*_*mes*_ (µm) = 115.451 + 6.717 *D*_*cell*_ (µm); R^2^ = 0.21, p < 0.0001; Figure S4). Hence, *g*_*ias*_ as computed here was a function of cell diameter and mesophyll porosity.

## Literature data for Figure 1

To supplement our 3D microCT dataset, data for *S*_*m*_ were collected from the literature. We calculated *SA*_*mes*_*/V*_*mes*_ from *S*_*m*_ as:

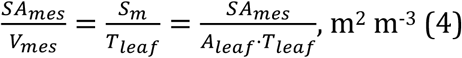

where *T*_*leaf*_ is leaf thickness and *A*_*leaf*_, leaf area. For this equation, we assumed that leaves were laminar and regular in shape, such that leaf volume could be approximated as a rectangular prism of *T*_*leaf*_ thickness and of a projected area equal to the leaf area. This assumption made this conversion only possible on laminar leaves.

## Statistical analysis

All analyses, simulations, and conductance computations were carried out in R 3.6.1^64^. Standardized major axes were computed using the smatr package^65^.

## Authors contributions

GTR, JME, and CRB planned the project, building from ideas of CKB and MJZ, and with contribution from CKB, MJZ, and MEG. GTR, JME, ABR, CRB, AJM, CKB, MJZ, and DT acquired microCT data. GTR and JME segmented the microCT images and extracted data from them. GTR and ABR planned the analysis, analyzed the data, and created the simulated dataset. KAS collected plant material and prepared samples for genome size analysis. DT contributed finite element modeling. GTR, ABR, KAS, and CRB wrote the manuscript, with contributions from all authors.

## Acknowledgements

We thank the University of California Botanical Garden (Berkeley, CA), the UC Davis Botanical Conservatory (Davis, CA), and the UC Davis Arboretum (Davis, CA) for plant material, the Paul Scherrer Institut, Villigen, Switzerland for provision of synchrotron radiation beamtime at beamline TOMCAT, and Klara Voggeneder for image processing. We thank the many who collected plant material on our behalf. The Advanced Light Source is supported by the Director, Office of Science, Office of Basic Energy Sciences, of the US Department of Energy under Contract no. DE-AC02-05CH11231. GTR was supported by a Katherine Esau Fellowship at UC Davis, and by the Austrian Science Fund (FWF), project M2245. This work was supported by US NSF grant DEB-1838327.

## Supplementary material

**Figure S1.**
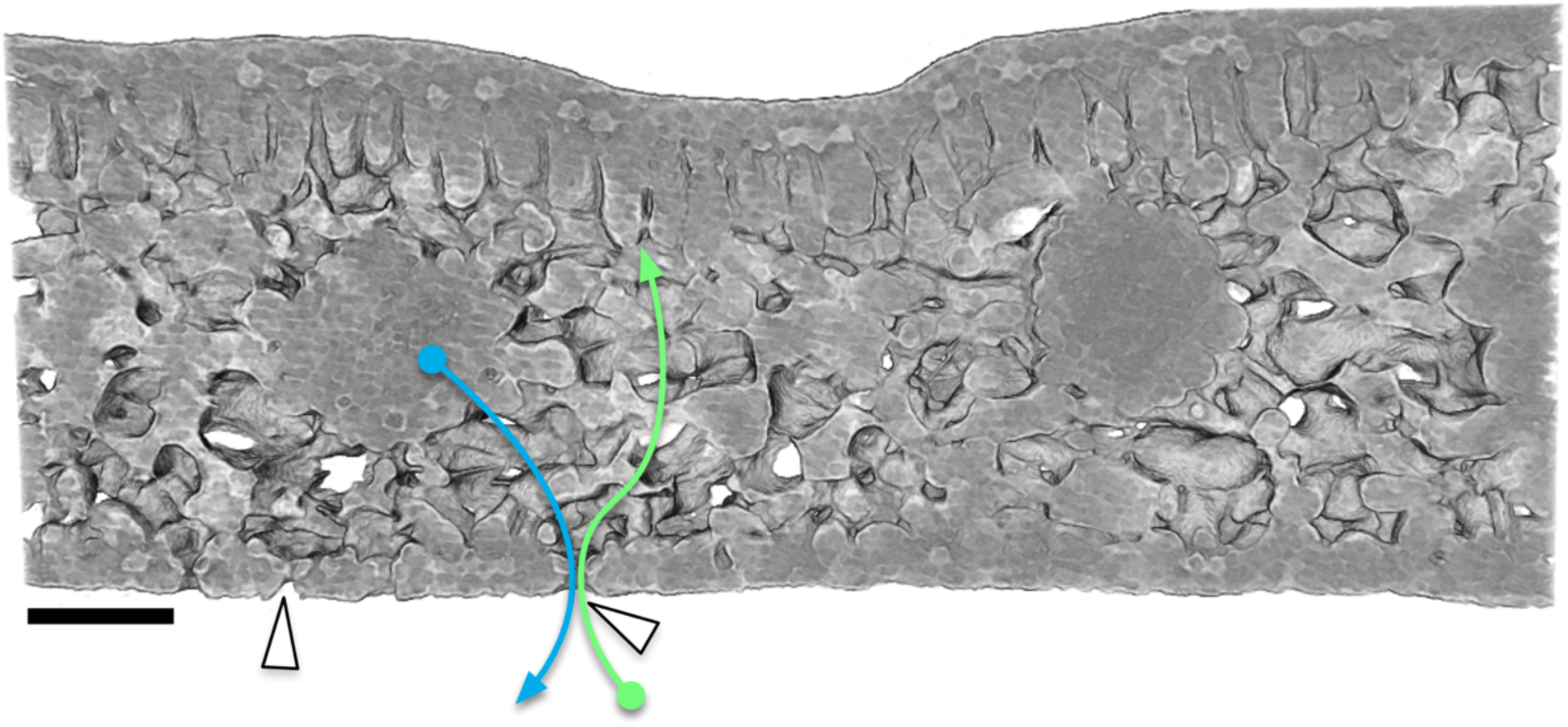
Transverse 3D X-ray microCT cross section through a *Agathis australis* leaf showing the complex 3D architecture of the leaf mesophyll. White arrows indicate locations of stomata on the abaxial surface. The blue arrow indicates one path a water molecule could travel from the vein out a stoma, and the green arrow indicates one path a CO_2_ molecule could travel through the stoma, through the spongy mesophyll, and into the palisade mesophyll near the adaxial surface.

**Figure S2.**
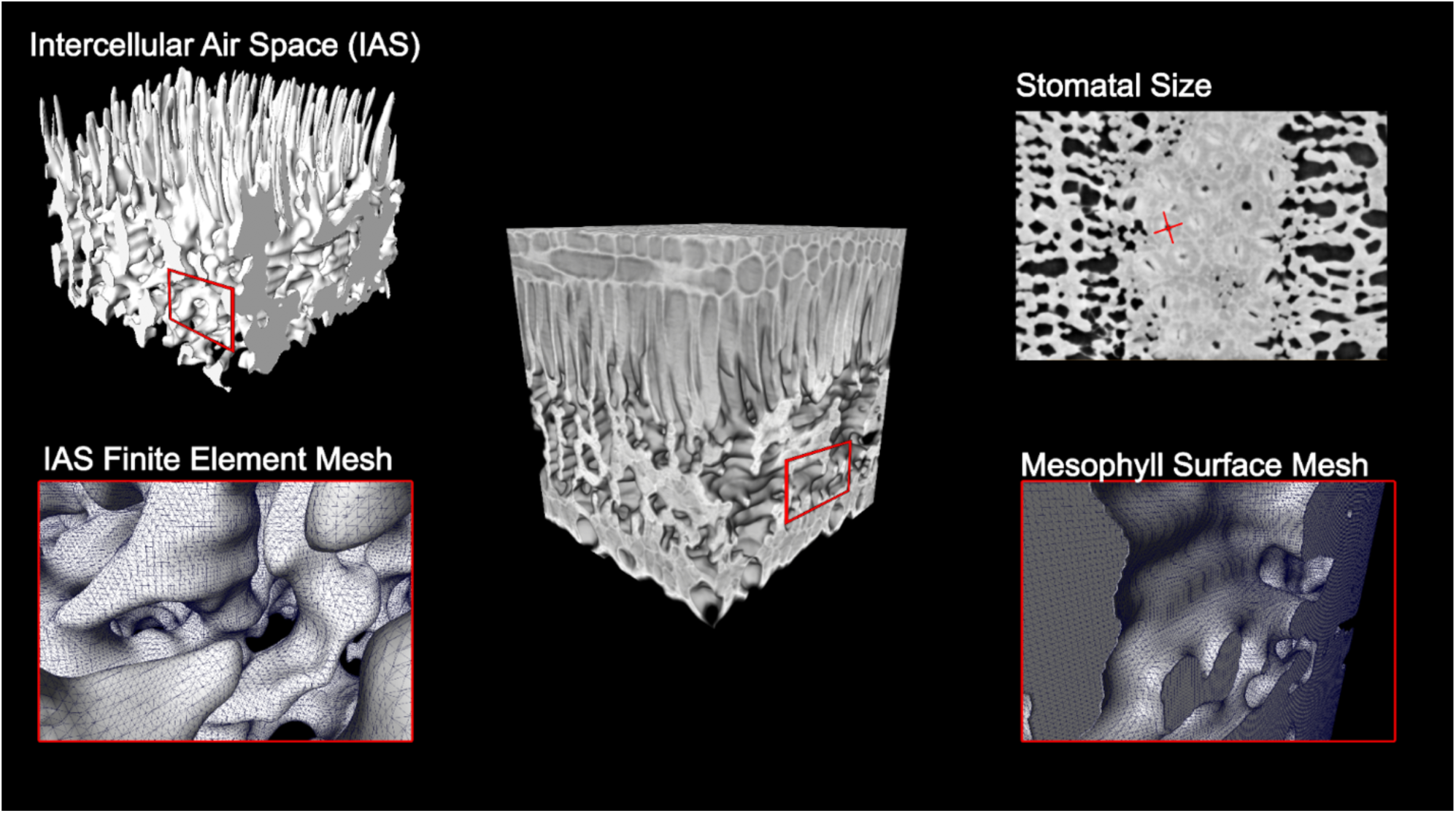
Two-dimensional and three-dimensional leaf traits can be measured on 3D microCT datasets. (center) A 3D volume rendering of a leaf. In 2D, traits such as guard cell length (top right) can be measured. In 3D, the intercellular airspace (top left) can be segmented and its surface (bottom left) measured by forming a finite element mesh. Similarly, the finite element mesh can be applied to the mesophyll cells (bottom right), allowing calculation of the surface area and volume.

**Figure S3.**
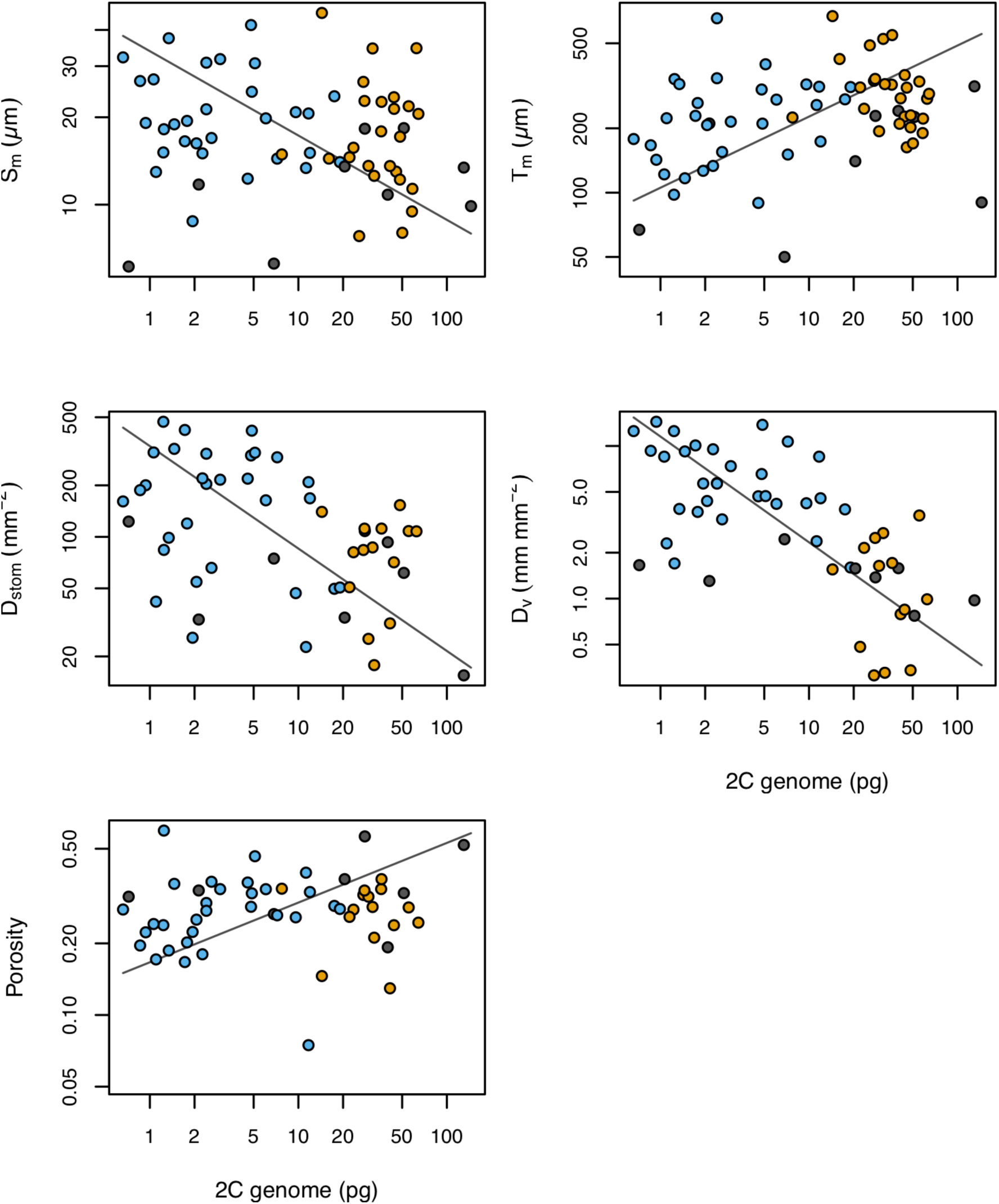
Relationship between 2C genome size (pg) and the surface area of mesophyll cells exposed to the intercellular airspace per leaf area (S_m_, R^2^=0.03, p=0.21), mesophyll thickness (T_m_, R^2^=0.09, p=0.015), stomatal density (D_stom_, R^2^=0.19, p=0.001), vein density (D_v_, R^2^=0.46, p<0.0001), and porosity (R^2^=0.01, p=0.38). Solid lines represent standard major axis regressions. Colored points represent angiosperms (blue), gymnosperms (orange), and ferns (gray).

**Table S1.**
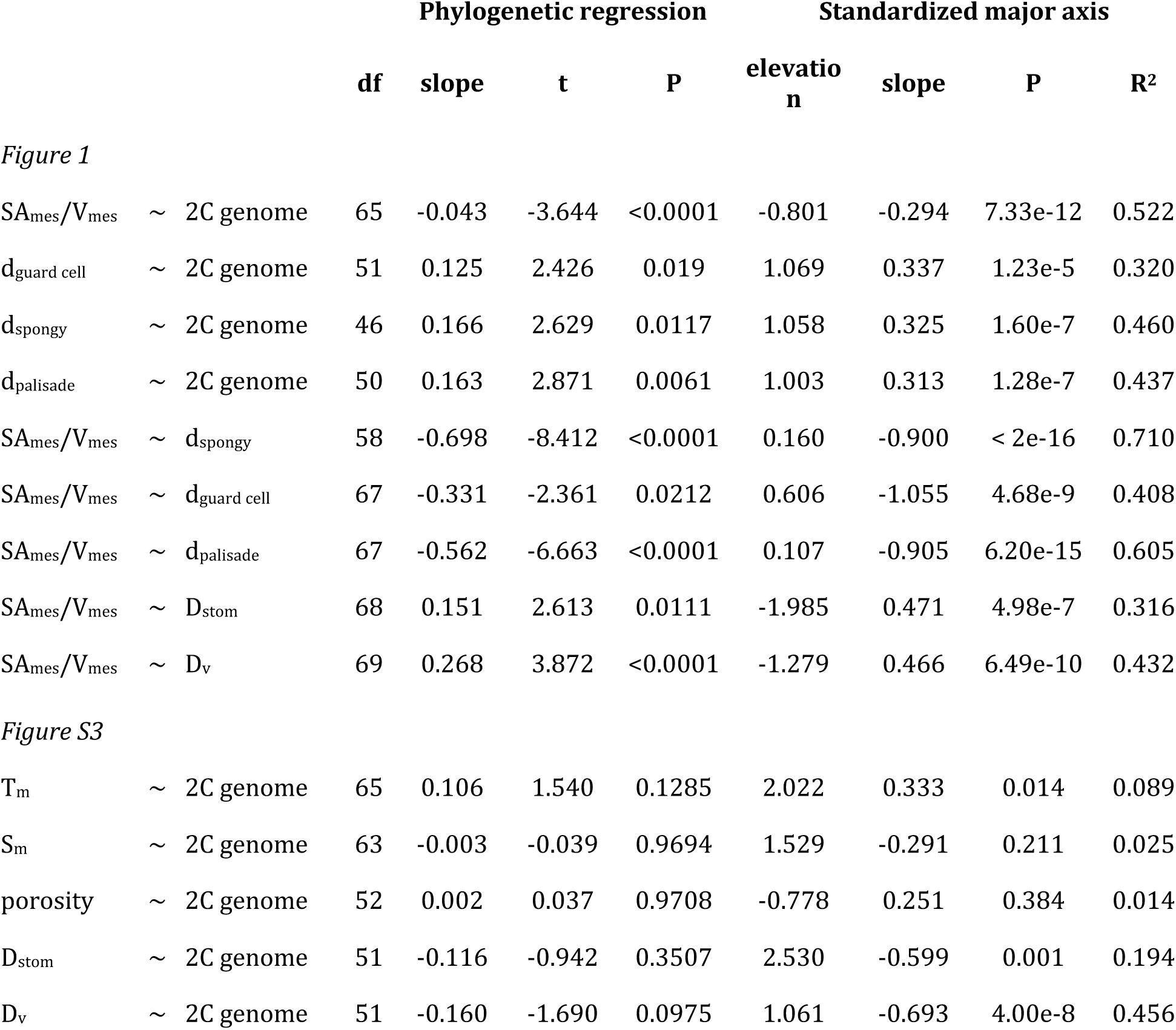
Phylogenetic least square regression and standardized major axis results for the relationship tested in Figures 1 and S3.

|  |  |  | Phylogenetic regression |  |  |  | Standardized major axis |  |  |  |
| --- | --- | --- | --- | --- | --- | --- | --- | --- | --- | --- |
|  |  |  | df | slope | t | P | elevation | slope | P | R<sup>2</sup> |
| *Figure 1* |  |  |  |  |  |  |  |  |  |  |
| $SA_{mes}/V_{mes}$ | ~ | 2C genome | 65 | -0.043 | -3.644 | <0.0001 | -0.801 | -0.294 | 7.33e-12 | 0.522 |
| $d_{guard\ cell}$ | ~ | 2C genome | 51 | 0.125 | 2.426 | 0.019 | 1.069 | 0.337 | 1.23e-5 | 0.320 |
| $d_{spongy}$ | ~ | 2C genome | 46 | 0.166 | 2.629 | 0.0117 | 1.058 | 0.325 | 1.60e-7 | 0.460 |
| $d_{palisade}$ | ~ | 2C genome | 50 | 0.163 | 2.871 | 0.0061 | 1.003 | 0.313 | 1.28e-7 | 0.437 |
| $SA_{mes}/V_{mes}$ | ~ | $d_{spongy}$ | 58 | -0.698 | -8.412 | <0.0001 | 0.160 | -0.900 | < 2e-16 | 0.710 |
| $SA_{mes}/V_{mes}$ | ~ | $d_{guard\ cell}$ | 67 | -0.331 | -2.361 | 0.0212 | 0.606 | -1.055 | 4.68e-9 | 0.408 |
| $SA_{mes}/V_{mes}$ | ~ | $d_{palisade}$ | 67 | -0.562 | -6.663 | <0.0001 | 0.107 | -0.905 | 6.20e-15 | 0.605 |
| $SA_{mes}/V_{mes}$ | ~ | $D_{stom}$ | 68 | 0.151 | 2.613 | 0.0111 | -1.985 | 0.471 | 4.98e-7 | 0.316 |
| $SA_{mes}/V_{mes}$ | ~ | $D_v$ | 69 | 0.268 | 3.872 | <0.0001 | -1.279 | 0.466 | 6.49e-10 | 0.432 |
| *Figure S3* |  |  |  |  |  |  |  |  |  |  |
| $T_m$ | ~ | 2C genome | 65 | 0.106 | 1.540 | 0.1285 | 2.022 | 0.333 | 0.014 | 0.089 |
| $S_m$ | ~ | 2C genome | 63 | -0.003 | -0.039 | 0.9694 | 1.529 | -0.291 | 0.211 | 0.025 |
| porosity | ~ | 2C genome | 52 | 0.002 | 0.037 | 0.9708 | -0.778 | 0.251 | 0.384 | 0.014 |
| $D_{stom}$ | ~ | 2C genome | 51 | -0.116 | -0.942 | 0.3507 | 2.530 | -0.599 | 0.001 | 0.194 |
| $D_v$ | ~ | 2C genome | 51 | -0.160 | -1.690 | 0.0975 | 1.061 | -0.693 | 4.00e-8 | 0.456 |

## References

1. Franks, P. J. & Beerling, D. J. Maximum leaf conductance driven by CO_2_ effects on stomatal size and density over geologic time. Proceedings of the National Academy of Sciences 106, 10343–10347 (2009).

2. Boer, H. J. de, Eppinga, M. B., Wassen, M. J. & Dekker, S. C. A critical transition in leaf evolution facilitated the cretaceous angiosperm revolution. Nature Communications 3, 1221 (2012).

3. Parkhurst, D. Internal leaf structure: A three-dimensional perspective. in On the economy of plant form and function: Proceedings of the sixth maria moors cabot symposium, evolutionary constraints on primary productivity, adaptive patterns of energy capture in plants, harvard forest, august 1983 (Cambridge [Cambridgeshire]: Cambridge University Press, c1986., 1986).

4. Evans, J. R., Kaldenhoff, R., Genty, B. & Terashima, I. Resistances along the CO_2_ diffusion pathway inside leaves. Journal of Experimental Botany 60, 2235–2248 (2009).

5. Earles, J. M. et al. Embracing 3D complexity in leaf carbon–water exchange. Trends in Plant Science 24, 15–24 (2019).

6. Simonin, K. A. & Roddy, A. B. Genome downsizing, physiological novelty, and the global dominance of flowering plants. PLoS Biology 16, e2003706 (2018).

7. Beerling, D. & Woodward, F. Changes in land plant function over the phanerozoic: Reconstructions based on the fossil record. Botanical Journal of the Linnean Society 124, 137–153 (1997).

8. Hetherington, A. & Woodward, F. The role of stomata in sensing and driving environmental change. Nature 424, 901–908 (2003).

9. Brodribb, T. J., Holbrook, N. M., Zwieniecki, M. A. & Palma, B. Leaf hydraulic capacity in ferns, conifers and angiosperms: Impacts on photosynthetic maxima. New Phytologist 165, 839–846 (2005).

10. Brodribb, T. J., Feild, T. S. & Jordan, G. J. Leaf maximum photosynthetic rate and venation are linked by hydraulics. Plant Physiology 144, 1890–1898 (2007).

11. Boyce, C. K., Brodribb, T. J., Feild, T. S. & Zwieniecki, M. A. Angiosperm leaf vein evolution was physiologically and environmentally transformative. Proceedings of the Royal Society B: Biological Sciences 276, 1771–1776 (2009).

12. Brodribb, T. J. & Feild, T. S. Leaf hydraulic evolution led a surge in leaf photosynthetic capacity during early angiosperm diversification. Ecology Letters 13, 175–183 (2010).

13. Feild, T. S. et al. Fossil evidence for cretaceous escalation in angiosperm leaf vein evolution. Proceedings of the National Academy of Sciences 108, 8363–8366 (2011).

14. Boyce, C. K. & Zwieniecki, M. A. Leaf fossil record suggests limited influence of atmospheric CO_2_ on terrestrial productivity prior to angiosperm evolution. Proceedings of the National Academy of Sciences 109, 10403–10408 (2012).

15. Brodribb, T. J., Jordan, G. J. & Carpenter, R. J. Unified changes in cell size permit coordinated leaf evolution. New Phytologist 199, 559–570 (2013).

16. John, G. P., Scoffoni, C. & Sack, L. Allometry of cells and tissues within leaves. American Journal of Botany 100, 1936–1948 (2013).

17. Feild, T. S. & Brodribb, T. J. Hydraulic tuning of vein cell microstructure in the evolution of angiosperm venation networks. New Phytologist 199, 720–726 (2013).

18. Baresch, A., Crifò, C. & Boyce, C. K. Competition for epidermal space in the evolution of leaves with high physiological rates. New Phytologist 221, 628–639 (2019).

19. Ši’mová, I. & Herben, T. Geometrical constraints in the scaling relationships between genome size, cell size and cell cycle length in herbaceous plants. Proceedings of the Royal Society B: Biological Sciences 279, 867–875 (2012).

20. Roddy, A. B. et al. The scaling of genome size and cell size limits maximum rates of photosynthesis with implications for ecological strategies. International Journal of Plant Sciences 181, 75–87 (2020).

21. Flexas, J. et al. Mesophyll conductance to CO_2_ and rubisco as targets for improving intrinsic water use efficiency in C3 plants. Plant, Cell & Environment 39, 965–982 (2016).

22. Lundgren, M. R. & Fleming, A. J. Cellular perspectives for improving mesophyll conductance. The Plant Journal doi:10.1111/tpj.14656.

23. Momayyezi, M., McKown, A. D., Bell, S. C. & Guy, R. D. Emerging roles for carbonic anhydrase in mesophyll conductance and photosynthesis. The Plant Journal doi: 10.1111/tpj.14638.

24. Tholen, D. et al. The chloroplast avoidance response decreases internal conductance to CO_2_diffusion in Arabidopsis thaliana leaves. Plant, Cell & Environment 31, 1688–1700 (2008).

25. Ren, T., Weraduwage, S. M. & Sharkey, T. D. Prospects for enhancing leaf photosynthetic capacity by manipulating mesophyll cell morphology. Journal of Experimental Botany 70, 1153–1165 (2019).

26. Théroux-Rancourt, G. et al. The bias of a two-dimensional view: Comparing two-dimensional and three-dimensional mesophyll surface area estimates using noninvasive imaging. New Phytologist 215, 1609–1622 (2017).

27. Earles, J. M. et al. Beyond porosity: 3D leaf intercellular airspace traits that impact mesophyll conductance. Plant Physiology 178, 148–162 (2018).

28. Smith, W. K., Vogelmann, T. C., DeLucia, E. H., Bell, D. T. & Shepherd, K. A. Leaf Form and Photosynthesis. BioScience 47, 785–793 (1997).

29. Oguchi, R., Onoda, Y., Terashima, I. & Tholen, D. Leaf anatomy and function. in The leaf: A platform for performing photosynthesis (eds. Adams III, W. W. & Terashima, I.) 97–139 (Springer International Publishing, 2018). doi:10.1007/978-3-319-93594-2_5.

30. Lundgren, M. R. et al. Mesophyll porosity is modulated by the presence of functional stomata. Nature Communications 10, 2825 (2019).

31. Tholen, D., Boom, C. & Zhu, X.-G. Opinion: Prospects for improving photosynthesis by altering leaf anatomy. Plant Science 197, 92–101 (2012).

32. Lehmeier, C. et al. Cell density and airspace patterning in the leaf can be manipulated to increase leaf photosynthetic capacity. The Plant Journal 92, 981–994 (2017).

33. Evans, J. & Vogelmann, T. C. Profiles of ^14^C fixation through spinach leaves in relation to light absorption and photosynthetic capacity. Plant, Cell & Environment 26, 547–560 (2003).

34. Borsuk, A. & Brodersen, C. R. The spatial distribution of chlorophyll in leaves. Plant Physiology 180, 1406–1417 (2019).

35. Feild, T. S. et al. Fossil evidence for low gas exchange capacities for early cretaceous angiosperm leaves. Paleobiology 37, 195–213 (2011).

36. West, G. B., Brown, J. H. & Enquist, B. J. A general model for the structure and allometry of plant vascular systems. Nature 400, 664–667 (1999).

37. Enquist, B. J. et al. Scaling metabolism from organisms to ecosystems. Nature 423, 639 (2003).

38. Kozłowski, J., Konarzewski, M. & Gawelczyk, A. T. Cell size as a link between noncoding DNA and metabolic rate scaling. Proceedings of the National Academy of Sciences 100, 14080–14085 (2003).

39. Price, C. A. et al. Testing the metabolic theory of ecology. Ecology Letters 15, 1465–1474 (2012).

40. Leitch, I. & Bennett, M. Genome downsizing in polyploid plants. Biological journal of the Linnean Society 82, 651–663 (2004).

41. Dodsworth, S., Chase, M. & Leitch, A. R. Is post-polyploidization diploidization the key to the evolutionary success of the angiosperms? Botanical Journal of the Linnean Society 180, 1–5 (2016).

42. Levin, D. A. Polyploidy and novelty in flowering plants. The American Naturalist 122, 1–25 (1983).

43. Doyle, J. J. & Coate, J. E. Polyploidy, the nucleotype, and novelty: The impact of genome doubling on the biology of the cell. International Journal of Plant Sciences 180, 1–52 (2019).

44. Baniaga, A., Marx, H. E., Arrigo, N. & Barker, M. S. Polyploid plants have faster rates of multivariate climatic niche evolution than their diploid relatives. Ecology Letters 23, (2020).

## Methods References

45. Gürsoy, D., De Carlo, F., Xiao, X. & Jacobsen, C. TomoPy: a framework for the analysis of synchrotron tomographic data. Journal of synchrotron radiation 21, 1188–1193 (2014).

46. Dowd, B. A. et al. Developments in synchrotron x-ray computed microtomography at the National Synchrotron Light Source. in SPIE’s international symposium on optical science, engineering, and instrumentation (ed. Bonse, U.) 224–236 (SPIE, 1999). doi:10.1117/12.363725.

47. Paganin, D., Mayo, S. C., Gureyev, T. E., Miller, P. R. & Wilkins, S. W. Simultaneous phase and amplitude extraction from a single defocused image of a homogeneous object. Journal of microscopy 206, 33–40 (2002).

48. Théroux-Rancourt, G. et al. Digitally deconstructing leaves in 3D using X-ray microcomputed tomography and machine learning. bioRxiv (2019) doi:10.1101/814954.

49. Schneider, C. A., Rasband, W. S. & Eliceiri, K. W. NIH Image to ImageJ: 25 years of image analysis. Nature methods 9, 671–675 (2012).

50. Doube, M. et al. BoneJ: Free and extensible bone image analysis in ImageJ. Bone 47, 1076–1079 (2010).

51. Bennett, M. D. & Leitch, I. J. Plant DNA C-values Database (release 6.0). http://data.kew.org/cvalues/.

52. Dolezel, J., Greilhuber, J. & Suda, J. Estimation of nuclear dna content in plants using flow cytometry. Nature Protocols 2, 2233–44 (2007).

53. Chamberlain, S. brranching: Fetch ‘phylogenies’ from many sources. (2019).

54. Magallon, S., Hilu, K. W. & Quandt, D. Land plant evolutionary timeline: Gene effects are secondary to fossil constraints in relaxed clock estimation of age and substitution rates. American Journal of Botany 100, 556–573 (2013).

55. Lu, Y., Ran, J.-H., Guo, D.-M., Yang, Z.-Y. & Wang, X.-Q. Phylogeny and divergence times of gymnosperms inferred from single-copy nuclear genes. PloS one 9, e107679 (2014).

56. Testo, W. & Sundue, M. A 4000-species dataset provides new insight into the evolution of ferns. Molecular Phylogenetics and Evolution 105, 200–211 (2016).

57. Webb, C. O., Ackerly, D. D. & Kembel, S. W. Phylocom: Software for the analysis of phylogenetic community structure and trait evolution. Bioinformatics 24, 2098–2100 (2008).

58. Pinheiro, J., Bates, D., DebRoy, S., Sarkar, D. & R Core Team. nlme: Linear and nonlinear mixed effects models. (2019).

59. Paradis, E. & Schliep, K. ape 5.0: An environment for modern phylogenetics and evolutionary analyses in R. Bioinformatics 35, 526–528 (2018).

60. Harwood, R. et al. Cell and chloroplast anatomical features are poorly estimated from 2D cross-sections. New Phytologist (2019) doi:10.1111/nph.16219.

61. Théroux-Rancourt, G., Voggeneder, K. & Tholen, D. Shape matters: The pitfalls of analyzing mesophyll anatomy. New Phytologist 1–4 (2019) doi:10.1111/nph.16219.

62. Theroux-Rancourt, G. & Gilbert, M. E. The light response of mesophyll conductance is controlled by structure across leaf profiles. Plant, Cell & Environment 40, 726–740 (2017).

63. Nobel, P. S. Physicochemical & environmental plant physiology. (Academic press, 2009).

64. R Core Team. R: A language and environment for statistical computing. (R Foundation for Statistical Computing, 2019).

65. Warton, D. I., Duursma, R. A., Falster, D. S. & Taskinen, S. smatr 3 - an R package for estimation and inference about allometric lines. Methods in Ecology and Evolution 3, 257–259 (2012).

